# CRISPR-Cas inhibits plasmid transfer and immunizes bacteria against antibiotic resistance acquisition in manure

**DOI:** 10.1101/2023.09.26.559507

**Authors:** Chahat Upreti, Pranav Kumar, Lisa M. Durso, Kelli L. Palmer

## Abstract

The horizontal transfer of antibiotic resistance genes among bacteria is a pressing global issue. The bacterial defense system CRISPR-Cas acts as a barrier to the spread of antibiotic resistance plasmids, and CRISPR-Cas-based antimicrobials can be effective to selectively deplete antibiotic-resistant bacteria. While significant surveillance efforts monitor the spread of antibiotic-resistant bacteria in the clinical context, a major, often overlooked aspect of the issue is resistance emergence in agriculture. Farm animals are commonly treated with antibiotics, and antibiotic resistance in agriculture is on the rise. Yet, CRISPR-Cas efficacy has not been investigated in this setting. Here, we evaluate the prevalence of CRISPR-Cas in agricultural *Enterococcus faecalis* strains and its anti-plasmid efficacy in an agricultural niche – manure. Analyzing 1,986 *E. faecalis* genomes from human and animal hosts, we show that the prevalence of CRISPR-Cas subtypes is similar between clinical and agricultural *E. faecalis* strains. Using plasmid conjugation assays, we found that CRISPR-Cas is a significant barrier against resistance plasmid transfer in manure. Finally, we used a CRISPR-based antimicrobial approach to cure resistant *E. faecalis* of erythromycin resistance, but this was limited by delivery efficiency of the CRISPR antimicrobial in manure. However, immunization of bacteria against resistance gene acquisition in manure was highly effective. Together, our results show that *E. faecalis* CRISPR-Cas is prevalent and effective in an agricultural setting and has the potential to be utilized for depleting antibiotic-resistant populations. Our work has broad implications for tackling antibiotic resistance in the increasingly relevant agricultural setting, in line with a One Health approach.

**Importance:** Antibiotic resistance is a growing global health crisis in human and veterinary medicine. Previous work has shown technologies based on CRISPR-Cas - a bacterial defense system - to be effective in tackling antibiotic resistance. Here we test if CRISPR-Cas is present and effective in agricultural niches, specifically in the ubiquitously present bacterium – *Enterococcus faecalis*. We show that CRISPR-Cas is both prevalent and functional in manure, and has the potential to be used to specifically kill bacteria carrying antibiotic resistance genes. This study demonstrates the utility of CRISPR-Cas based strategies for control of antibiotic resistance in agricultural settings.

## 1. Introduction

A widely prevalent adaptive immunity system possessed by bacteria and archaea is Clustered Regularly Interspaced Short Palindromic Repeats (CRISPR)-Cas ^1,2^. Although it has now been programmed as a tool to edit genomes of a large number of organisms^3–6^, in its native environment, it confers bacteria and archaea defense against mobile genetic elements (MGE) like viruses and plasmids^1,2,7,8^. Upon entry of the MGE, a portion of its DNA, known as a protospacer, is integrated into the host genome in the form of a spacer^8^. Later, the spacer is transcribed (called crRNA) and loaded on Cas-encoded nuclease(s) to surveille for future MGE entries. If a region of an incoming MGE is complementary to the crRNA, the MGE is cleaved by the nuclease^9–11^. Since its discovery, a large majority of studies on the CRISPR-Cas mechanism of action have focused on its antiviral activity. However approximately 10% of CRISPR protospacers identified in bacteria and archaea target plasmids, suggesting that CRISPR-Cas naturally provides defense against invading plasmids as well^12^. Indeed, CRISPR-Cas has been shown to act as a natural barrier against conjugative plasmid transfer^13,14^.

A highly relevant clinical organism in the study of CRISPR-Cas function is *Enterococcus faecalis. E. faecalis* is a Gram-positive bacterium routinely isolated from a large range of hosts including animals, humans and environmental samples such as plants, sand, and water^15,16,17^. It is an opportunistic pathogen and causes diseases including urinary tract infections, infective endocarditis and bacteremia^18–21^. Owing in part to its significant repertoire of intrinsic antibiotic resistance genes and its ability to efficiently acquire resistance genes from external sources, *E. faecalis* is one of the leading causes of hospital-acquired infections^22,23^. *E. faecalis* also colonizes several food animals and strains from these sources have been reported to carry both antibiotic resistance and virulence genes ^24,25^. Further, due to the generalist nature of *E. faecalis*, transmission of resistant *E. faecalis* from animals to humans (zoonosis) can occur. Indeed, there have been several cases of probable zoonotic transmission leading to major outbreaks in the past^26,27^, and enterococci, including *E. faecalis*, are routinely monitored in retail meat as part of the National Antimicrobial Resistance Monitoring System^28^.

Some *E. faecalis* strains encode Type II CRISPR-Cas systems which employ a crRNA-Cas9 nuclease-tracrRNA complex to surveille and cleave MGEs^29–31^. *E. faecalis* has two functional CRISPR-Cas loci – CRISPR1-Cas and CRISPR3-Cas. Both have a CRISPR array consisting of 36 bp repeats interspersed with 30 bp spacers. Both loci have their own set of *cas* genes and protospacer adjacent motif (PAM) sequences, suggesting distinct modes of action and targets. Almost every *E. faecalis* strain examined so far has also been found to have another CRISPR locus called CRISPR2, however it lacks *cas* genes and is thus not independently functional^32^. Since plasmid-mediated transfer of antibiotic resistance genes is a major contributor to resistance transmission in *E. faecalis*^23,33^, it is important to test if CRISPR-Cas could act as a barrier against this process, especially in conditions where zoonotic transmission may occur. Additionally, the prevalence and function of CRISPR-Cas in strains of animal origin should be investigated, as most focus has been on human-origin strains. Some recent studies have looked at *E. faecalis* samples isolated from wastewater treatment plants, animal feces and soil. However, these studies have either small sample sizes, low source diversity or unclear origin of strains^34–36^.

Previous work has shown *E. faecalis* CRISPR-Cas systems confer defense against plasmids and can be programmed as antimicrobials against specific antibiotic-resistant strains, both *in vitro* and *in vivo* in the mouse gut ^14,37^. During in vitro mating experiments, CRISPR-Cas acted as a moderately effective barrier against plasmid transfer, while in vivo its efficiency was strikingly robust^14^, suggesting that the conjugation environment impacts CRISPR-Cas function. Given the importance of agriculturally relevant niches in *E. faecalis* biology, it is crucial to test how effective CRISPR-Cas is in this setting. A key niche of this setting is manure. Animal manure, which is frequently used as fertilizers, are a known reservoir of antibiotic resistance gene-encoding pathogenic bacteria including *E. faecalis.* Thus they are potential hubs for zoonotic transmission^38–40^, as well as potential critical control points. Since previous work in *E. faecalis* has demonstrated the surrounding environmental plays a major role in CRISPR-Cas efficacy^14^, discerning its efficacy in manure could be of major significance.

In this study, we first identified the prevalence of CRISPR-Cas in a large cohort of agricultural *E. faecalis* strains. Next, we evaluated CRISPR-Cas efficacy in sterilized manure using previously established model *E. faecalis* strains. Finally, we tested the effectiveness of CRISPR-antimicrobials designed to eliminate antibiotic resistance genes from *E. faecalis* in manure.

## 2. Results

### CRISPR-Cas prevalence in agricultural *Enterococcus faecalis* is similar to that in human-derived strains

Before testing the efficacy of CRISPR-Cas in a model agricultural setting, we wanted to determine how prevalent CRISPR-Cas is in agriculturally-derived *E. faecalis* and how it compares to its prevalence in strains derived from humans. For this, we downloaded all *E. faecalis* genomes from NCBI Refseq available as of September 1, 2021, along with genomes from a recent large-scale OneHealth investigation of Enterococci^25^. We segregated the genomes based on source of isolation – human or animal (the term animal here is an umbrella term for non-human sources like food animals, birds, pets etc.). Together, these comprised 1587 genomes sourced from humans (79.91%) and 399 sourced from animals (20.09%) (Fig 1A), for a total of 1,986 genomes. To identify CRISPR-Cas loci in these genomes, we performed a BLAST search using the *cas9* sequences of the *E. faecalis* CRISPR1-Cas and CRISPR3-Cas loci as queries, and then searched for bona fide CRISPR arrays within the set of genomes containing *cas9*. We focused on CRISPR1-Cas and CRISPR3-Cas for this analysis since these are the only two independently functional CRISPR arrays in *E. faecalis*. We found that 27.57% of animal-derived strains possessed CRISPR1-Cas, and 9.52% strains possessed CRISPR3-Cas. This was similar to their prevalence in human-derived strains – 29.36% for CRISPR1-Cas and 8.5% for CRISPR3-Cas (Fig 1B). Next, we investigated the CRISPR arrays of these strains to determine whether and to what extent the targets of CRISPR-Cas systems in these two groups differ. For this, we extracted the CRISPR-Cas arrays corresponding to the established *E. faecalis* CRISPR loci from the genomes, and collated the spacers across them separately for animal and human derived genomes. For assigning targets to these spacers, we created a database of plasmids isolated from *E. faecalis, E. faecium and E. hirae*, as well as the NCBI viral genome database. With respect to plasmid targets, we found 95 targets for the set of spacers from animal derived *E. faecalis*, and 114 targets for spacers derived from human sources. The Venn diagram in Fig 1C shows that all targets of animal derived strains were also in the list of targets of human derived strains. With respect to viral targets, we found 134 targets for animal derived strains, and 114 targets for human derived strains. Again, a large majority of the targets (100) overlapped between the two groups of targets, suggesting major similarity in CRISPR-Cas targets across the two environments. This suggests that CRISPR-Cas prevalence is consistent across human and animal niches along with major similarities in their targets, with the caveat that our study is limited to the *E. faecalis* strains that have been selected for sequencing by other groups, which may have biases (i.e. prioritization of antibiotic-resistant strains for sequencing).

**FIG 1.**
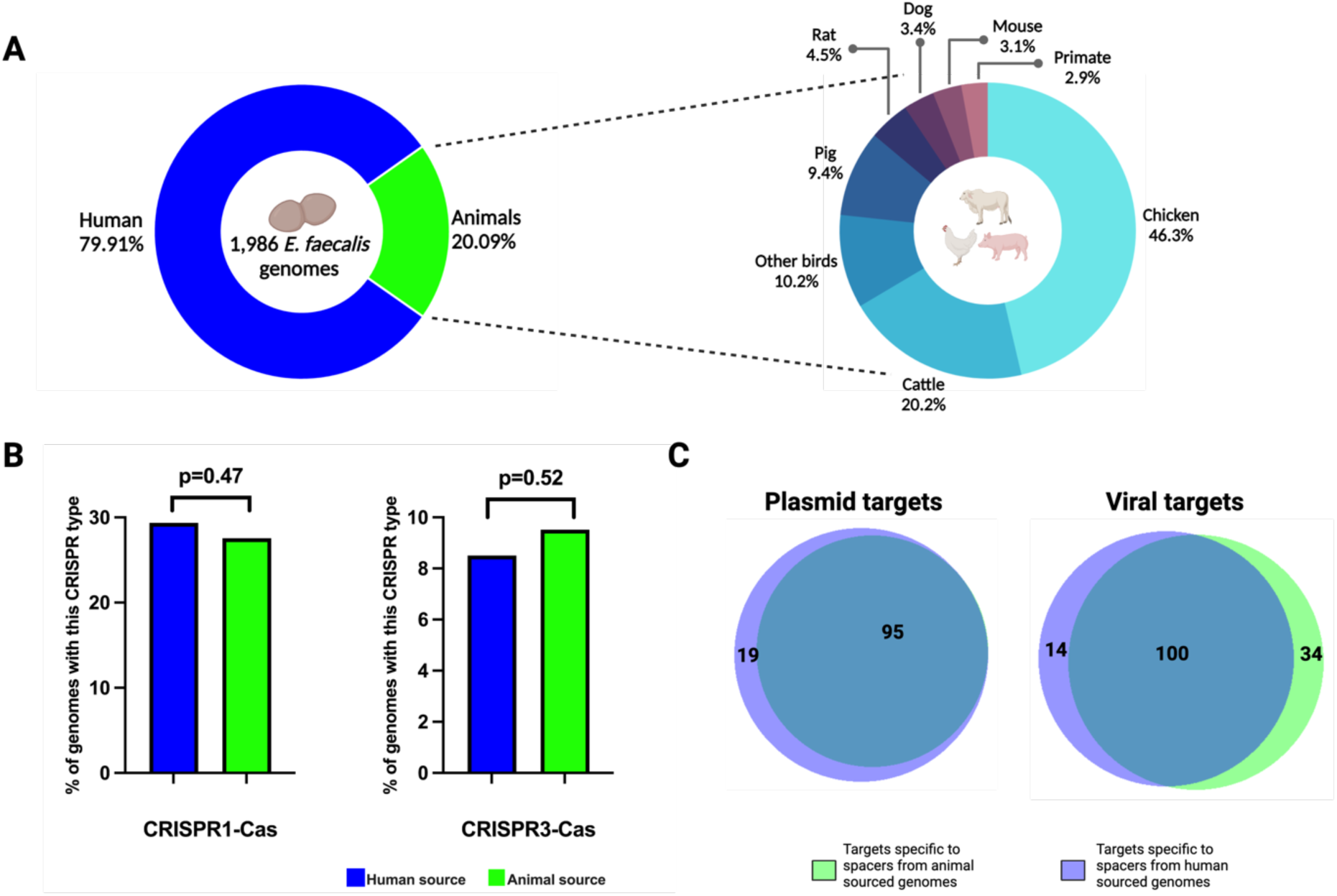
*E. faecalis* strains studied and their CRISPR-Cas abundances and targets. (A) shows the distribution of *E. faecalis* strains from human and animal sources analyzed in this study. (B) shows relative abundances of CRISPR-Cas types in *E. faecalis* strains from animal and human sources. Percentages were calculated as the number of strains in each source category (animal/human) containing a particular CRISPR-Cas type (CRISPR1-Cas or CRISPR3-Cas) divided by the total number of strains in that source category. p-values were calculated using the Z test for equality of two percentages using independent samples. (C) shows Venn diagrams for plasmid and viral targets of spacers derived from animal and human sourced *E. faecalis*.

### CRISPR-Cas blocks plasmid transfer in manure, and manure-specific effects on CRISPR defense are observed

We next tested the hypothesis that *E. faecalis* CRISPR-Cas is effective in an agricultural niche (manure). We tested the efficacy of CRISPR-Cas in blocking the dissemination of pAM714 from *E. faecalis* OG1SSp to *E. faecalis* T11RF (Fig 2). For this, we performed conjugation assays in three conditions - solid brain heart infusion (BHI) agar, liquid BHI broth, and liquified autoclaved dairy manure (Fig 3). Two different recipients were used, T11RF with a functional CRISPR-Cas system, and T11RFΔ*cas9* which lacks functional CRISPR-Cas defense against pAM714 (Table 1)^13,14^. For these experiments, donor and recipient cells were mixed together and either spread on BHI agar to form a biofilm, or inoculated into liquid BHI, or manure. After 18 hours incubation, agar plate biofilms were harvested and resuspended in buffer, and liquid cultures (liquid BHI and manure) were sampled as well. These were then plated for determination of colony forming units (CFU) of donors, recipients, and transconjugant cells. This timepoint is referred to as passage 0.

**FIG 2.**
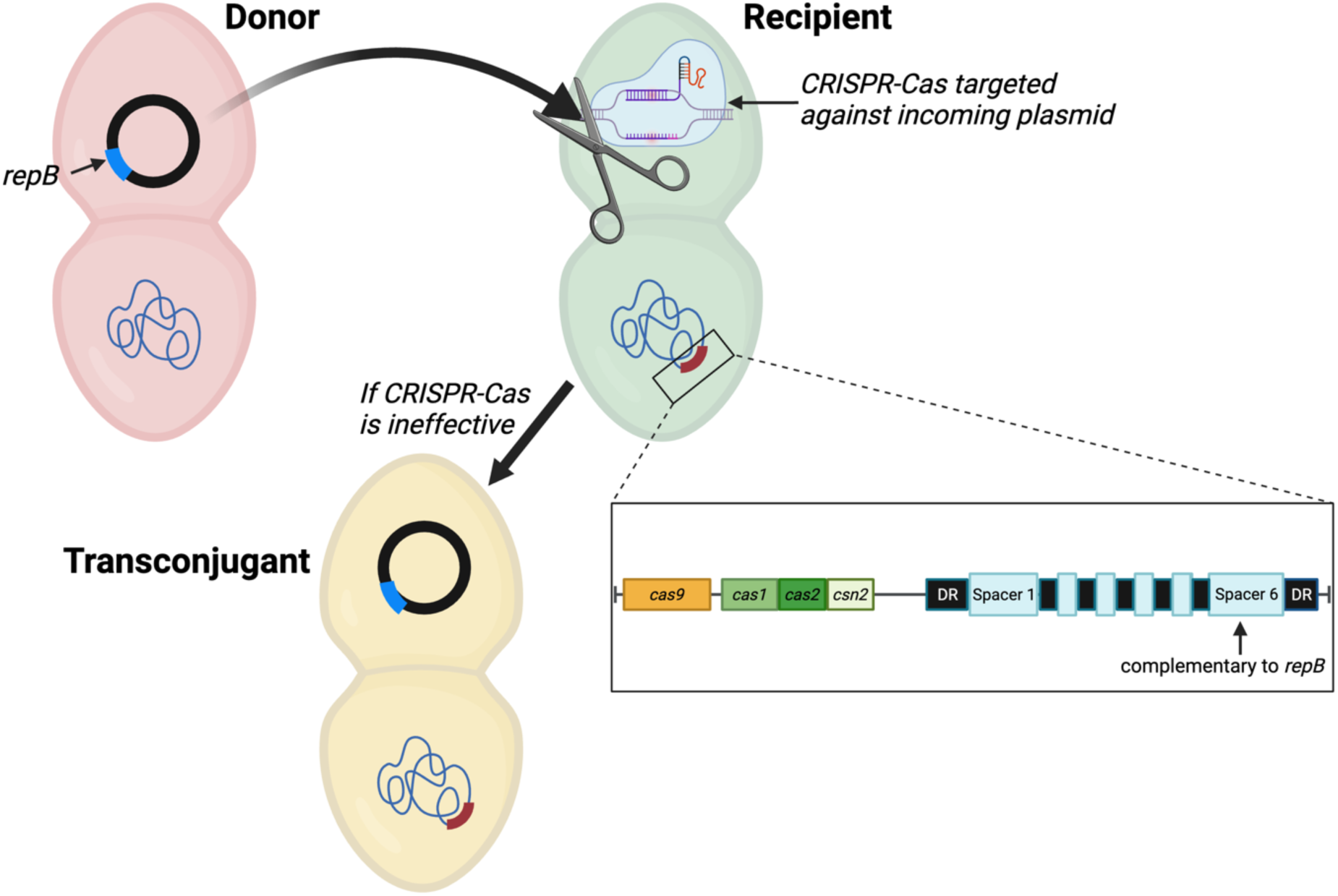
A schematic representation of the concept. The donor strain OG1SSp contains a pheromone-responsive plasmid which harbors a *repB* gene (blue). The recipient strain T11RF encodes a Type-II CRISPR-Cas system (red) containing a spacer with perfect complementarity to *repB*. If CRISPR-Cas is effective in a particular niche, it will block the entry of the incoming plasmid. If not, the plasmid enters the recipient, which is then termed a transconjugant.

**FIG 3.**
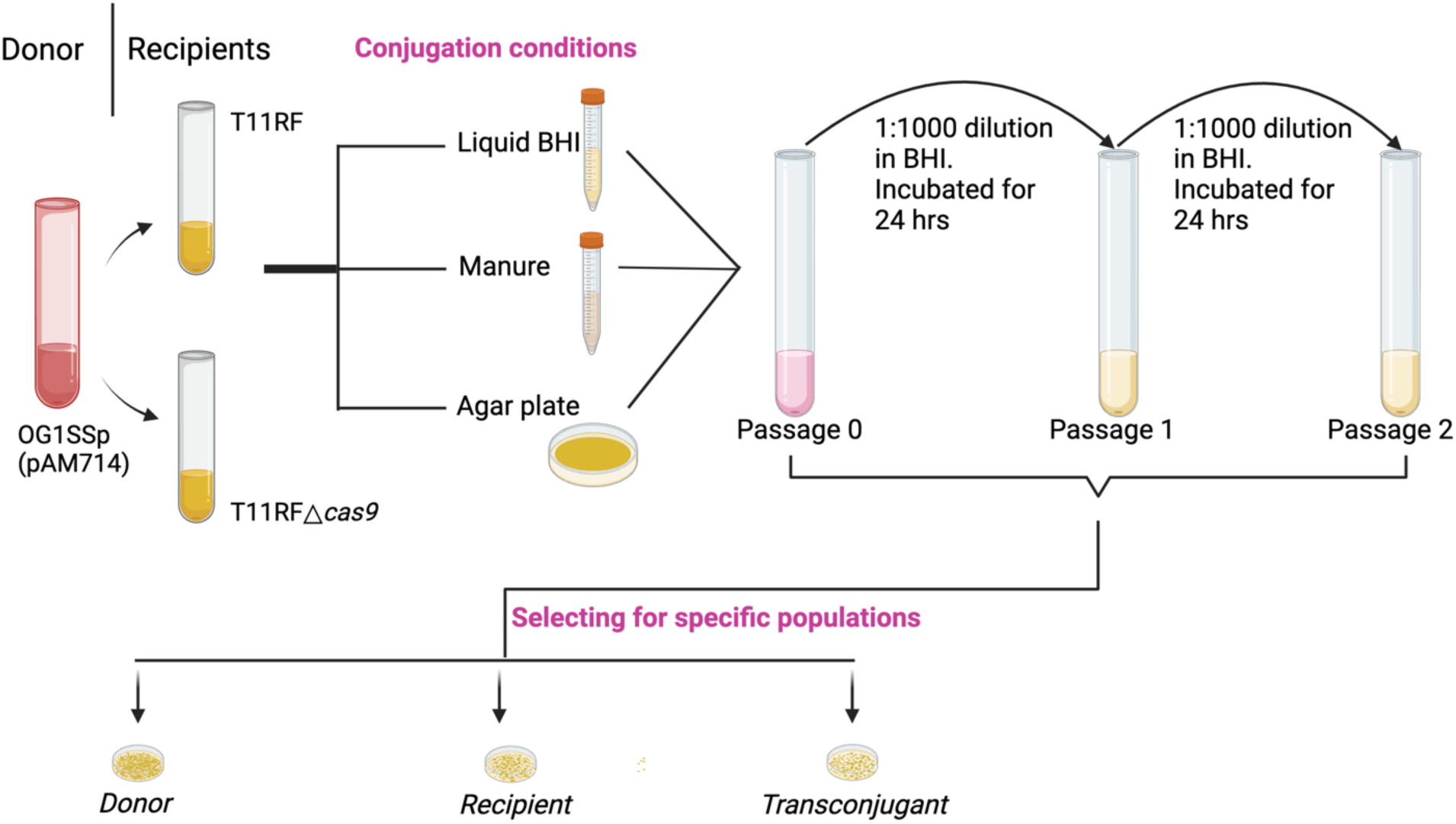
A schematic of the experimental design. The donor strain (OG1SSp(pAM714)) was mixed with two versions of the recipient (T11RF, T11RFΔ*cas9*) in three conditions – liquid BHI, manure and BHI agar plate. After 18 hours, the mixture (Passage 0) was plated to quantify donor (spectinomycin, streptomycin, and erythromycin selection), recipient + transconjugants (rifampin and fusidic acid selection) and transconjugant (rifampin, fusidic acid, and erythromycin selection) populations. At the same time, Passage 0 was also diluted into fresh liquid BHI (1:1000 dilution) and incubated for 24 hours to give Passage 1. The Passage 1 culture was similarly used to generate Passage 2.

**Table 1.**
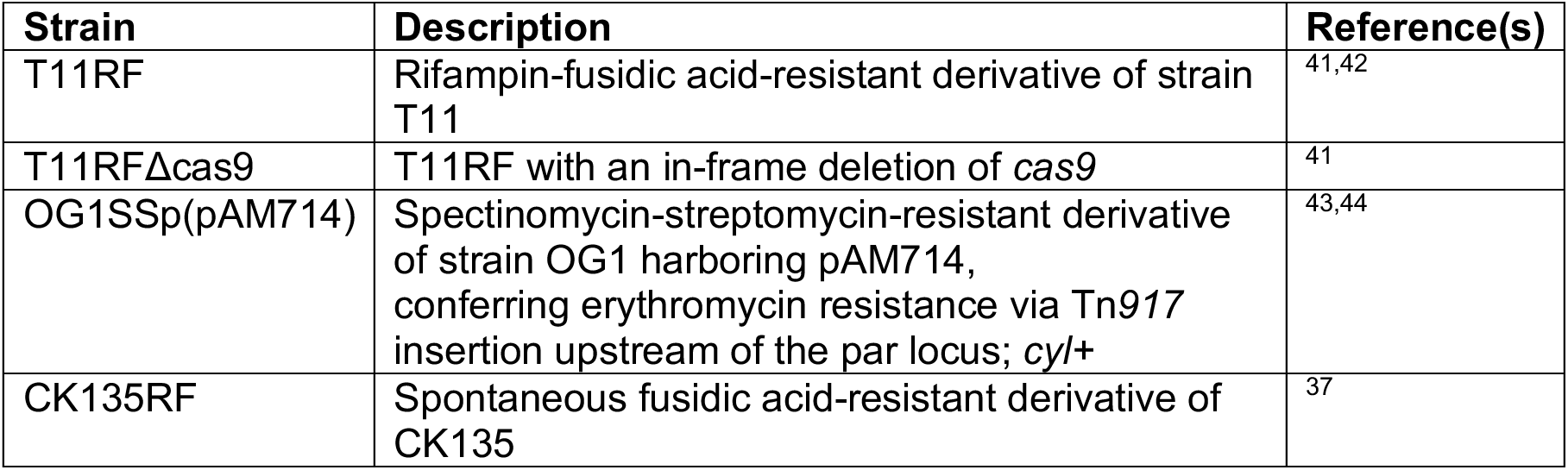
*E. faecalis* strains used in this study.

In passage 0, we found transconjugants in the range of 10^4^–10^7^ CFU/mL across the three environments and both recipients, suggesting widespread plasmid transfer across all 6 combinations (Fig 4). However, there was a significant difference in pAM714 conjugation frequency (number of transconjugants divided by the number of donors) on BHI agar in the absence vs presence of *cas9*, with a higher conjugation frequency in the absence of functional CRISPR-Cas (Fig 5a). This was expected since it has been reported previously, demonstrating that CRISPR-Cas is a moderately effective barrier against plasmid transfer on solid agar^14^. This effect was also seen in liquid BHI medium, and importantly, also in manure, although the difference in conjugation frequency in manure for T11RF versus T11RFΔ*cas9* recipients was less than the other conditions at this time point.

**FIG 4.**
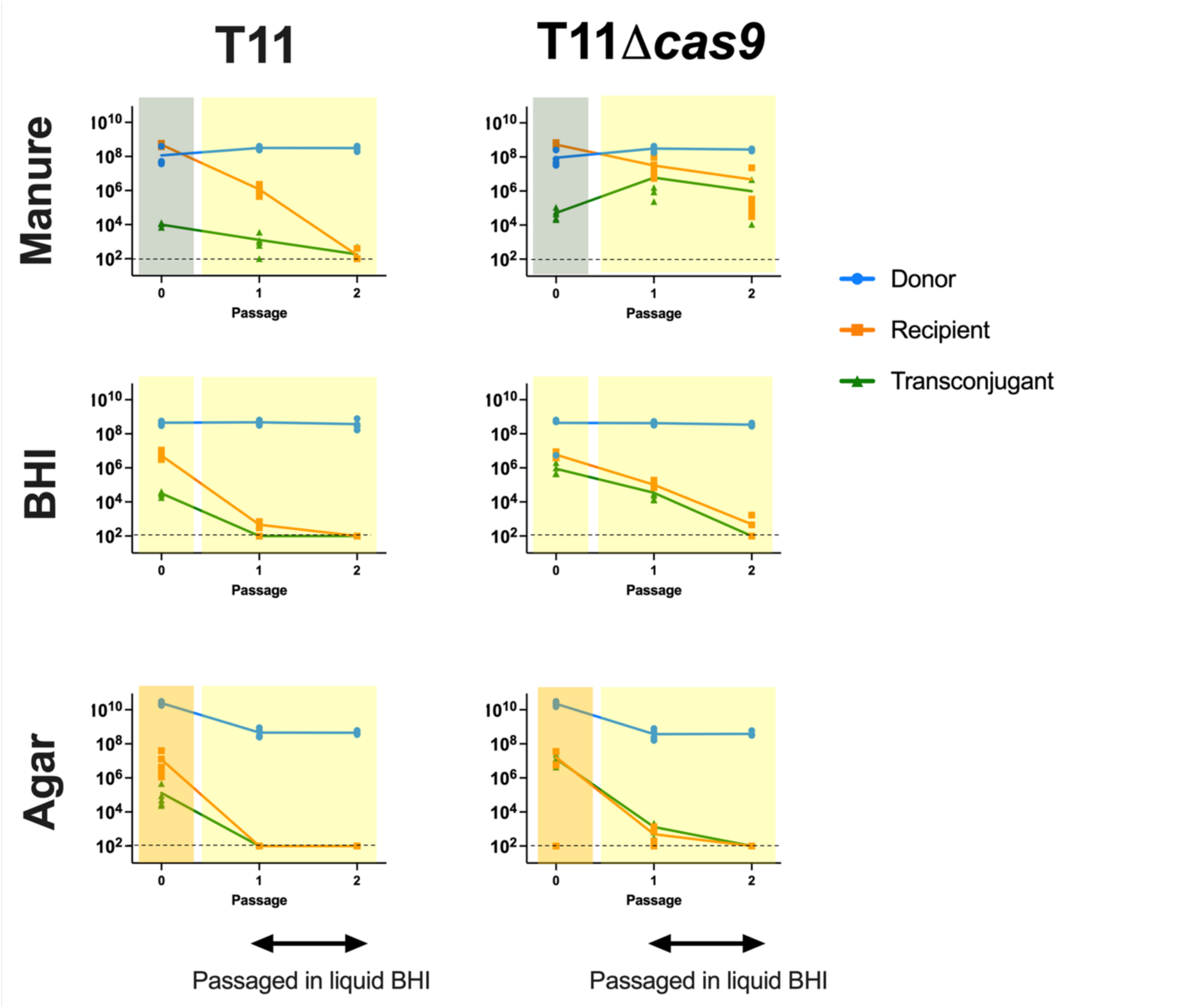
Bacterial populations across time and niche. Each graph shows the number of donors, recipients, and transconjugants in different environments. The conjugation experiments shown on the left had T11RF as the recipient and those on right had T11RFΔ*cas9*. Colors in the background represent the condition in which the bacteria were growing: manure (dark green), liquid BHI (yellow) and solid agar plate (light orange). Dashed lines represent a CFU/ml count of 10^2^ which is the limit of detection.

**FIG 5.**
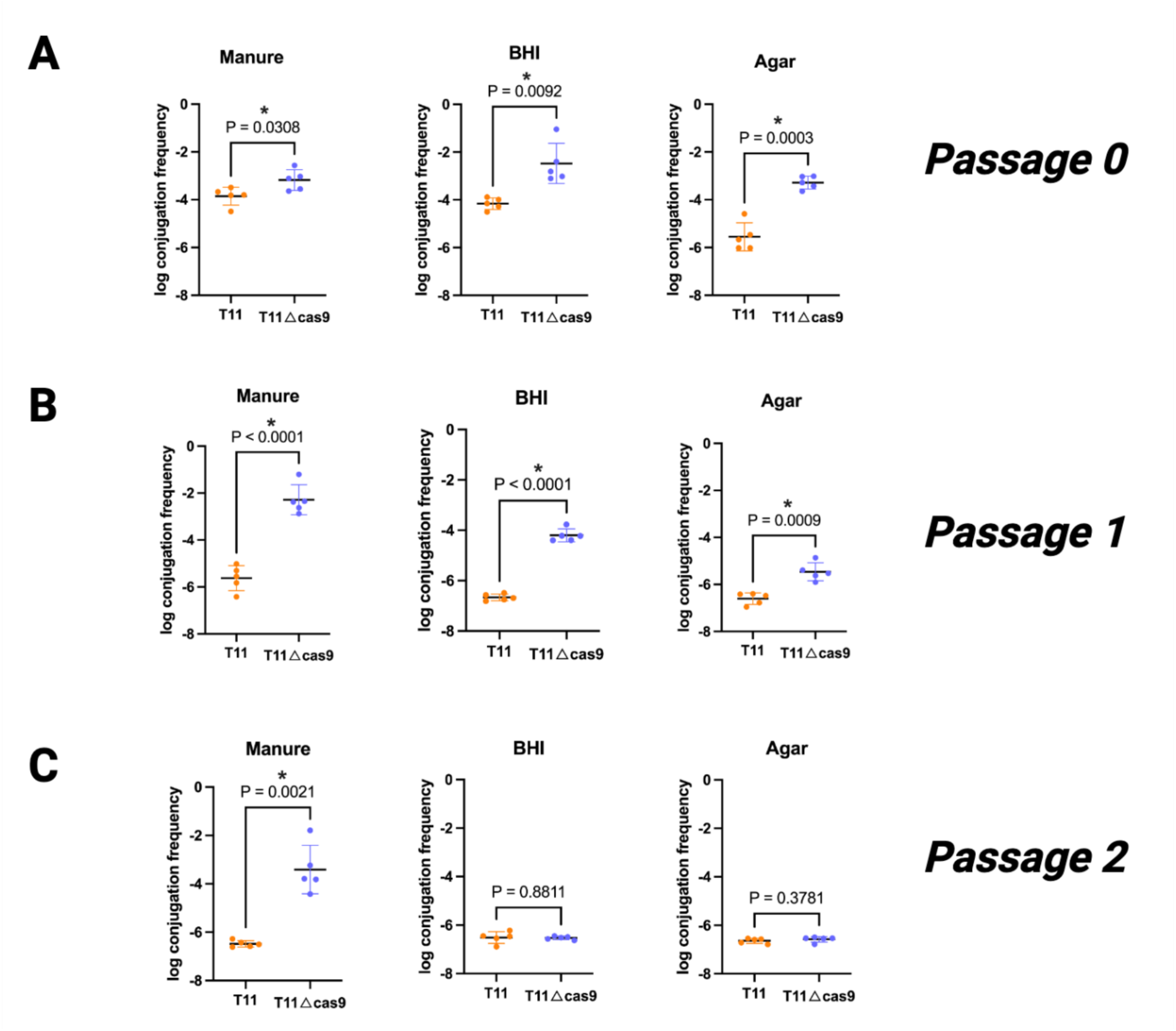
CRISPR-Cas conjugation frequency under different conditions. Each point represents log of conjugation frequency (calculated as number of transconjugants divided by number of donors) corresponding to each of the three initial conjugation conditions (manure, BHI broth, BHI agar). A represents Passage 0 (right after initial conjugation reaction); B and C represent subsequent overnight passages - Passage 1 and Passage 2. Student’s t-test was used to calculate p-values after determining that log conjugation frequency values followed a normal distribution.

We also performed two successive passages of these initial conjugation mixtures. For passage 1, we diluted the passage 0 conjugation mixture 1:1000 into fresh BHI broth and incubated it at 37°C for 24 hours, followed by plating on selective agars. Transferring the conjugation mixture from manure exposure to liquid BHI was a way to model transition between two nutritional environments. Here, we found the conjugation frequencies in the presence of CRISPR-Cas as opposed to its absence, continued to be significantly lower across all three conditions (p<0.0001, p<0.0001, p=0.0009 for manure, agar and BHI respectively) (Fig 5b). This demonstrates that CRISPR-Cas continued to be effective at blocking plasmid transfer in this sample set.

Next, passage 1 conjugation mixtures were diluted and incubated as described above to obtain passage 2 cultures. By the end of passage 2, the transconjugant populations in agar and liquid BHI media were almost completely absent (limit of detection of 10^2^ CFU/mL), irrespective of CRISPR-Cas. As a result, no differences in conjugation frequencies across the two conditions were observed (Fig 5c). Surprisingly, for conjugation mixtures that originated from manure, there was a sizeable transconjugant population following passage 2, but only in the absence of functional CRISPR-Cas (Fig 4<u>, top right</u>). As a result, the conjugation frequency in the absence of CRISPR-Cas was significantly higher than in the presence of CRISPR-Cas (Fig 5c). These results suggest a manure-specific effect on either CRISPR-Cas function and/or the competitiveness of T11RF Δ*cas9*(pAM714) transconjugants.

### CRISPR antimicrobials appear effective in manure, but are limited by low conjugation frequency

Having shown that CRISPR-Cas can function in an agriculturally-relevant *in vitro* setting, we next wanted to see if CRISPR-Cas-based antimicrobials can be used to selectively deplete antibiotic-resistant *E. faecalis* in that niche. Previous studies have tested their antimicrobial efficacy in solid agar and the murine gastrointestinal tract^37^. Solid agar is a popular choice for conjugation experiments due to the simplicity of its make-up and experimental convenience, but for CRISPR-based antimicrobials to be widely used, they need to be tested in settings that better mimic the *in situ* environment. Manure is one such environmental niche, since it plays an important role in the emergence and dissemination of resistant strains in agricultural settings. For this experiment, we used a previously designed CRISPR antimicrobial referred to as pKH88^37^. pKH88 is a derivative of the pheromone-responsive plasmid, pPD1, and has been modified to express *cas9* and spacers targeting either *tetM* (for tetracycline resistance) or *ermB* (for erythromycin resistance). For our experiments, we delivered pKH88[sp-*tetM*] or pKH88[sp- *ermB*] from the *E. faecalis* strain CK135RF (donor strain), which is a derivative of the strain OG1. Our target (recipient) strain was OG1SSp(pAM714), which encodes *ermB* on pAM714. Figure 6 shows a schematic diagram of our approach. In this scenario, our goals were to deliver pKH88[sp-*ermB*] to OG1SSp(pAM714), and then either (a) kill or inhibit the recipients’ growth by Cas9-mediated DNA damage, or (b) promote the loss of pAM714 via Cas9-mediated DNA damage. We expected no effects of delivering pKH88[sp-*tetM*] to OG1SSp(pAM714), as OG1SSp(pAM714) does not encode *tetM*; pKH88[sp-*tetM*] is our control.

**FIG 6.**
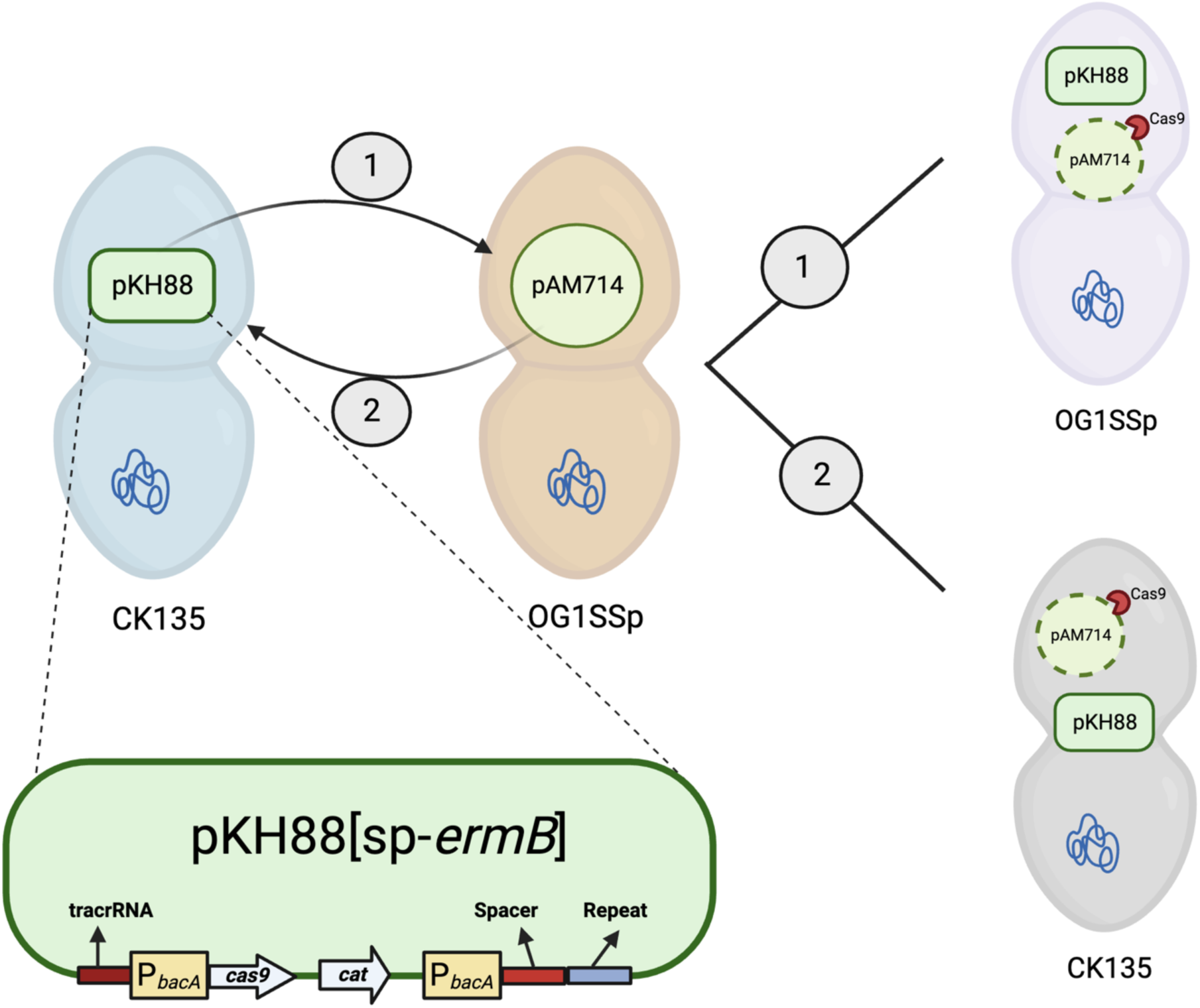
A schematic representation of the CRISPR antimicrobial. The strain CK135RF contains the pheromone-responsive plasmid pKH88, which encodes *cas9* and a CRISPR guide RNA under a constitutively active *bacA* promoter. The guide RNA consists of a spacer that targets either the *ermB* gene or the *tetM* gene, and a repeat sequence. The *cat* gene, encoding chloramphenicol resistance, is inserted in the plasmid only for tracking purposes. The strain OG1SSp contains the plasmid pAM714 which harbors the *ermB* gene. In scenario 1, the CRISPR antimicrobial pKH88[sp-*ermB*] enters OG1SSp and cleaves pAM714. In scenario 2, OG1SSp(pAM714) donates pAM714 to CK135RF(pKH88[sp-*ermB*]) and the CRISPR antimicrobial cleaves the incoming plasmid.

With this setup, we tested the hypothesis that the pKH88[sp-*ermB*] CRISPR antimicrobial plasmid could selectively deplete erythromycin resistance from *E. faecalis* populations cultured in manure. CRISPR antimicrobial donor strains and OG1SSp(pAM714) recipients were mixed and plated on solid agar or inoculated into liquid BHI or manure, and passages 0, 1 and 2 were assessed for donor, recipient, and transconjugant CFU/mL as described above. We observed a significant decrease in the erythromycin resistant population by passage 2 for agar plate conjugations (Fig 7a), in line with previously published results^37^. However, a similar decrease in the erythromycin-resistant population was not observed for liquid BHI or manure conditions in any of the passages (Fig 7a). To investigate a possible cause for this, we measured the transconjugant population in each case. We found that the transconjugant populations were substantially lower in liquid BHI and manure across all passages (10^2^ (limit of detection) to 10^7^), as compared to agar (10^9^ to 10^10^) (Fig 7b). A possible reason for this could be the absence of a solid surface for the bacteria to form a biofilm in the case of liquid BHI and manure, which promotes conjugation^45^. However, in both conditions and across all three passages (with the exception of passage 0 in liquid BHI), there were fewer transconjugants in the case of the experimental antimicrobial pKH88-*ermB* as compared to the control pKH88[sp-*tetM*] (Fig 7b), and this reduction was significant in passages 1 and 2 in manure. Assuming similar plasmid transfer levels for both pKH88[sp-*ermB*] and pKH88[sp-*tetM*], this could be explained as the action of the antimicrobial: OG1SSp(pAM714) recipients that received pKH88[sp-*ermB*] had their pAM714 cleaved due to CRISPR-Cas targeting and hence were not detected in transconjugant-selective plates. This suggests that the antimicrobial was effective in manure when conjugation did occur.

**FIG 7.**
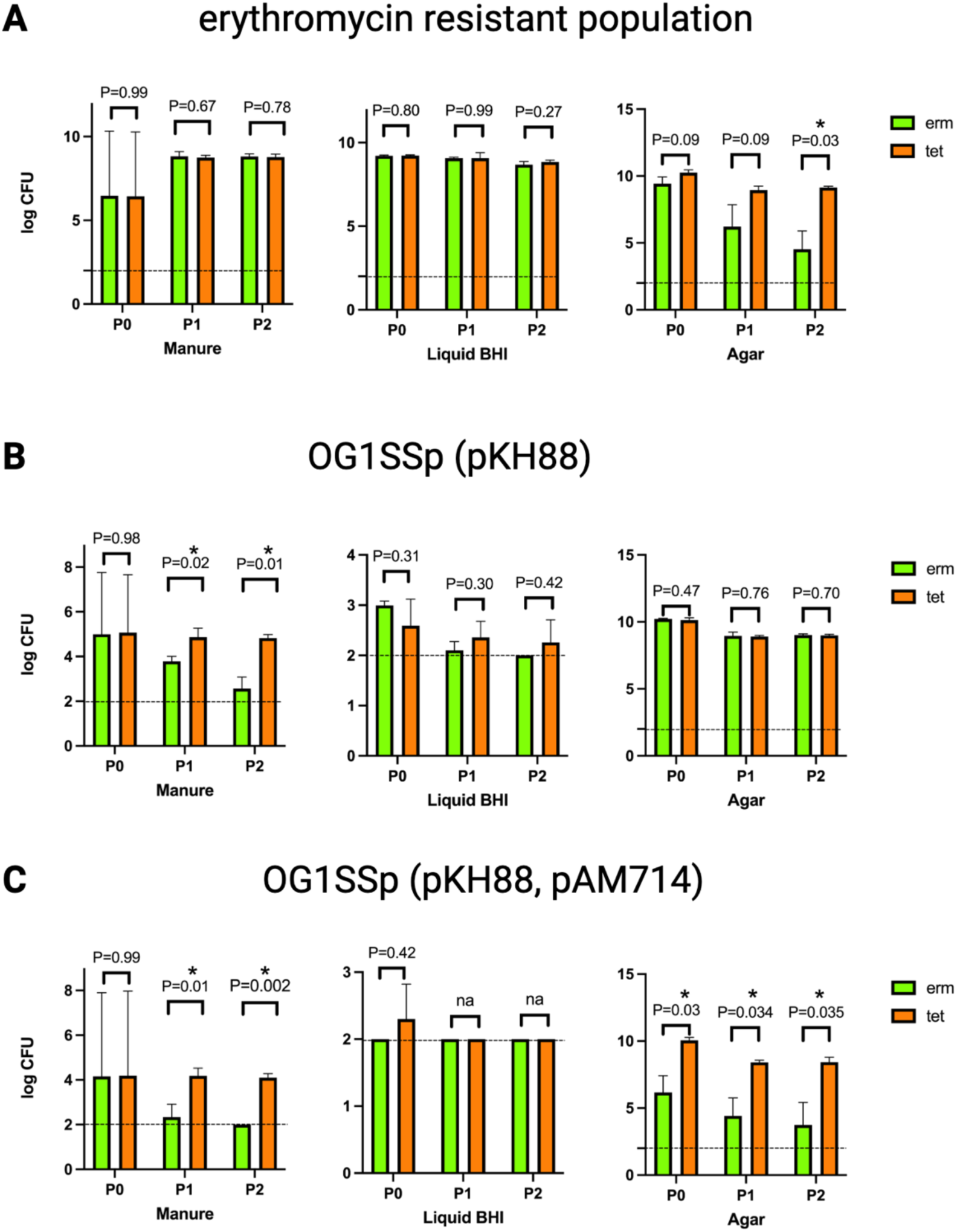
Evaluating CRISPR antimicrobial efficiency. (A) shows the number of total erythromycin-resistant bacteria in the mixture, across the three conditions, 3 passages and with the control (tet, in brown) or experimental (erm, in green) CRISPR antimicrobial. A lower number on this graph signifies depletion of the erythromycin-resistant population. (B) shows the number of transconjugants i.e. the recipient (OG1SSp) cells that have accepted the antimicrobial plasmid pKH88. (C) shows the number of OG1SSp(pKH88) transconjugants that still maintain the pAM714 plasmid. All values were log transformed and t-test with Welch’s correction was used to generate p-values. ‘na’ represents cases where values in both groups were the same and hence a t-test could not be performed. Dashed lines represent a CFU/ml of 10^2^ which is the limit of detection.

To further test the efficacy of the antimicrobial, we looked at the subset of the transconjugant population that was erythromycin-resistant. These would be OG1SSp(pAM714) recipient cells that received the pKH88[sp-*ermB*] or [sp-*tetM*] antimicrobial plasmid and also maintained erythromycin resistance. We found that in almost every passage for both manure and agar, this cell population was significantly lower in the case of the experimental antimicrobial pKH88[sp-*ermB*] as compared to the control pKH88[sp-*tetM*] (Fig 7c). This again points to the effectiveness of the antimicrobial.

Finally, we tested if the presence of the CRISPR antimicrobial could immunize the donor CK135RF(pKH88) bacteria against acquiring the *ermB* gene from OG1SSp(pAM714). For this we calculated the number of CK135RF(pKH88) bacteria that were erythromycin-resistant, which could be the result of pAM714 transferring from OG1SSp to CK135RF. We observed a stark difference in this population between the experimental and control antimicrobial. The number of erythromycin-resistant CK135RF(pKH88) was significantly lower in the case of the *erm* targeting antimicrobial as compared to the one that targeted the *tet* gene (Fig 8). This shows that the pKH88[sp-*ermB*] CRISPR antimicrobial immunized its host CK135RF against acquisition of erythromycin resistance. This effect was seen in both manure and agar.

**FIG 8.**
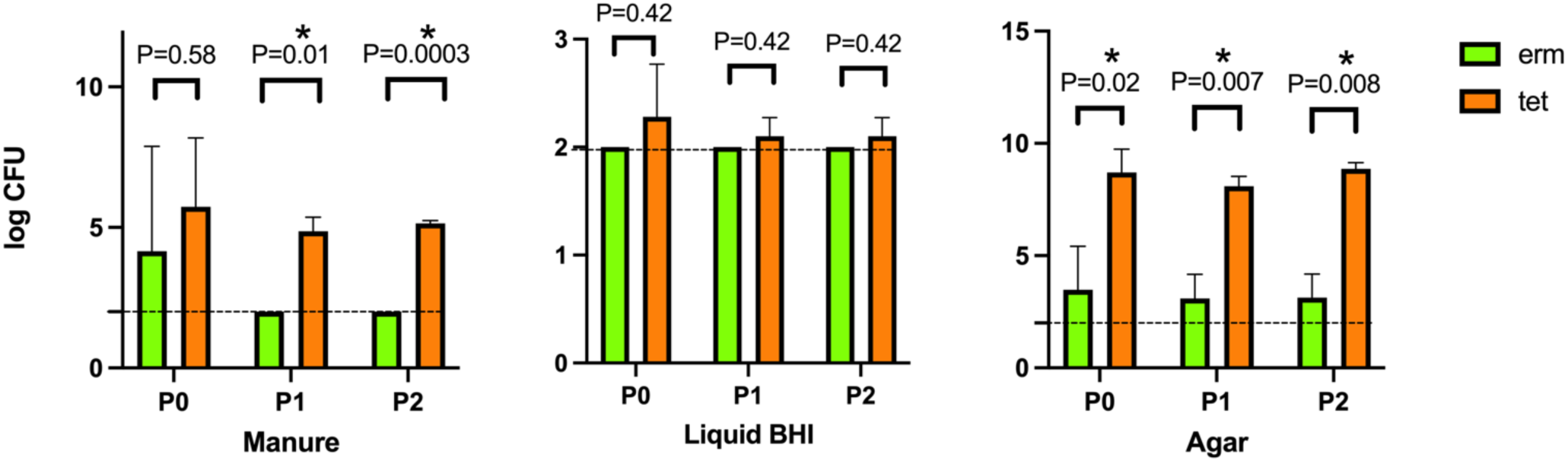
Immunization of bacteria against antibiotic resistance acquisition in manure. Bars represent number of CK135RF cells that have become erythromycin-resistant, likely via accepting pAM714 from OG1SSp, in addition to carrying pKH88. A lower number represents immunization via the antimicrobial plasmid pKH88. All values were log transformed and t-test with Welch’s correction was used to generate p-values. Dashed lines represent a CFU/ml of 10^2^ which is the limit of detection.

## 3. Materials and Methods

### Bacteria and reagents used

*E. faecalis* strains were grown in brain heart infusion (BHI) broth or on agar plates. Antibiotic concentrations used were as follows: rifampin, 50 μg/ml; fusidic acid, 25 μg/ml; spectinomycin, 500 μg/ml; streptomycin, 500 μg/ml; erythromycin, 50 μg/ml; tetracycline, 10 μg/ml; chloramphenicol, 15 μg/ml.

### Database collection and CRISPR-Cas prevalence analysis

NCBI Genome was used to download *E. faecalis* genomes publicly available as of September 1, 2021. Based on the host (source of isolation), the downloaded genomes were categorized as originating from human or animal sources. Genomes with hosts that did not match either of these categories were excluded. A recent study provided a significant number of additional genomes to our study that were not part of NCBI Genome (Refseq)^25^. Genomes were subject to filtering based on genome size to eliminate anomalies. The average genome size was calculated, and genomes with sizes less than half or greater than 1.5 times of this average size were discarded (following Refseq recommendation). Strains were queried on BLAST against the *cas9* sequences of CRISPR1-Cas (GCF_000172575.2) and CRISPR3-Cas (GCF_000157475.1) to discover strains with potentially active CRISPR-Cas (percent identity >= 98.5%). CRISPRCasFinder was used to identify CRISPR arrays in these genomes, and the known repeat sequences of CRISPR 1,2,3 were used to filter bona-fide CRISPR arrays from the result^30,46^.

### CRISPR target analysis

To investigate CRISPR-Cas targets, we used CRISPR-CasFinder to extract CRISPR arrays for all genomes. Next, using the direct repeats of CRISPR1-Cas, CRISPR2 and CRISPR3-Cas^30^, we tested whether the arrays extracted were indeed from one of these known loci by matching the known direct repeats with that of the extracted array, allowing for a Levenshtein distance of up to 3. For the set of bona fide arrays in both animal and human groups, we combined all the spacers across the three CRISPR-Cas subtypes. Next, we created a plasmid database by using PLSDB to gather plasmids isolated from *E. faecalis, E. faecium* and *E. hirae*, along with plasmids extracted from recently published *E. faecalis* strains isolated from gut and urine^47–49^. The viral database was created by downloading the viral part of the nr/nt database from NCBI, comprising ∼45,000 sequences. We then performed a BLAST search for the spacer sets against both these databases, with word size parameter as 8, and hits with at least 93% sequence identity and 93% query coverage were accepted as bona fide protospacers. The code for this analysis is available at https://github.com/chahatupreti/Agro-CRISPR.

### Manure microcosm

Dairy cattle manure was obtained from a settling basin in Kimberley, Idaho, and autoclaved to destroy resident flora. The solid manure was filtered through a cheesecloth to remove large fecal solids, and 1 mL of the resultant liquid manure was used per conjugation reaction.

### Conjugation experiments

Donors and recipients were grown overnight (from freezer stocks) in liquid BHI medium, followed by a 1:10 subculture. The subsample was statically incubated at 37°C for 1.5 hours after which time optical density (OD) measurement at 600 nm was taken to ensure similar ODs for both donor and recipient. For each microcosm, 100 µl of the donor culture and 900 µl of a recipient culture were centrifuged, 90% of the supernatant was removed, and then cells were resuspended. Three experimental conditions were used to study conjugation outcomes – solid brain heart infusion (BHI) agar plate, liquid BHI and liquid manure. Conjugation reactions were incubated for 18 hours at 37°C. After this, the conjugation mixture was collected, serial dilutions were performed and plated on agar plates with appropriate antibiotic selection. Where appropriate, the recipient population was calculated by subtracting CFU/mL of transconjugants from that of recipients+transconjugants. Plates were incubated at 37°C for 72 hours followed by counting the number of colony forming units/mL (CFU/mL). Conjugation frequency was calculated as 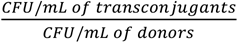. For multi-passage experiments, a 1:1000 dilution subculture was prepared and allowed to incubate in liquid BHI for 24 hours before another set of plating. For CRISPR antimicrobial experiments, the pheromone cAD1 (500 ng/mL) was added to the conjugation mixture to increase conjugation.

## 4. Discussion

In this study, we assessed the prevalence and function of CRISPR-Cas in agriculturally relevant niches. We discovered that the prevalence of both CRISPR1-Cas and CRISPR3-Cas (the two independently functional CRISPR-Cas types in *E. faecalis*) was similar when comparing strains of animal origin to those of human origin. These two CRISPR-Cas systems differ in terms of their protospacer-adjacent motif (PAM) used by the Cas nuclease to cleave the protospacer in the invading genome, along with differences in the corresponding *cas* genes^13^. The strong similarity in their prevalence suggests that mechanistic differences between the two subtypes do not have an impact on their distributions in *E. faecalis* from diverse hosts. Investigating the targets of the CRISPR systems showed that there was also remarkable similarity in the plasmid and viral targets of these systems across the two environments.

We also assessed how effective CRISPR-Cas is an agricultural niche, using dairy manure in a microcosm design. Previous studies reported that CRISPR-Cas efficacy varies with the environment it is tested in, with differences observed *in vitro* versus *in vivo*^14^. Our data show *E. faecalis* CRISPR-Cas is able to block plasmid transfer following exposure to manure. The effect of manure on CRISPR function is observable even after two post-manure passages in BHI medium, something that was not observed in the control BHI broth and agar microcosms. We speculate that even though the manure was autoclaved and most of its solids removed before using, it still contained factors that could be altering *E. faecalis* physiology. Specifically, T11RFΔ*cas9* remains at high levels in BHI-passaged cultures post-manure exposure (Figure 4). This is not observed for T11RF post-manure, or any of the cultures originating in BHI. Teasing apart the mechanism by which this happens and the relative contributions of manure, CRISPR-Cas, and pAM714 would be the next step. One approach could be to simplify the complex manure environment – manure could be fractionated by size and we could study if components larger or smaller than a particular size can fully capture the effect seen here. We could also target specific biomolecules in manure – lipids, proteins, DNA – by digesting them and seeing its effect on CRISPR-Cas efficacy. Another approach would be to test a different recipient strain to see how manure affects its CRISPR functioning and its growth/stability in culture. Since V583 is genetically highly similar to T11 (except for the mobile elements) while also being a highly drug-resistant strain, it would be a good choice of recipient to test this effect.

Overall, the microcosm studies provided clear evidence of *E. faecalis* CRISPR-Cas functioning in an agricultural (manure) environment and highlighted future challenges to be considered in experiments exploring potential field applications of the concept. First, our manure model is highly simplified – we autoclaved it to remove all resident microbiota and then filtered and strained it to remove most of the solids. The former was done to avoid complications that could arise from the resident microbiota affecting CRISPR-Cas function, and the latter to help standardize quantification of manure amounts. The solids, if present, could provide additional surface for the bacteria to form biofilms on, which would promote conjugation and hence, possibly, enhance the effect of CRISPR antimicrobials. Although we saw CRISPR antimicrobials have some effect, we were unable to bring up the conjugation rates to a sufficient extent to robustly test the effectiveness of the antimicrobial. It is possible that the large size of our antimicrobial plasmid was the reason for its low conjugation rate; engineering the antimicrobial using a smaller plasmid could potentially help improve its conjugation rate and performance in agricultural niches. Secondly, the experiments could be performed with the resident microbiota intact by using a combination of antibiotic-selective and species-selective plates. Thirdly, after letting the bacteria conjugate in the three respective microcosms, we subsequently passaged the conjugation mixture in liquid BHI in all three conditions, as a way to model transition across different nutritive environments which can happen as strains are transmitted between different hosts and niches. An alternative approach could be to passage each of the three arms in the same microcosm that they were exposed to during the initial period of conjugation (solid agar, liquid BHI and manure).

Our work has several important implications both in terms of basic CRISPR biology and tackling antibiotic resistance. Our analysis looking at CRISPR-Cas prevalence and targets in *E. faecalis* is perhaps the largest of its kind, and the insight that the distribution of CRISPR-Cas subtypes and targets is invariant across animal and human sources points to a lack of CRISPR-mediated host colonization preference. The *E. faecalis* microcosm studies offer a proof-of-concept on the potential for CRISPR-Cas to limit horizonal gene transfer in dairy manures. Finally, while conjugation efficiency limited the ability of our CRISPR antimicrobial to spread in manure to antibiotic-resistant target cells, we did successfully immunize an *E. faecalis* strain against antibiotic resistance acquisition, which was observed to be more robust in manure compared to liquid BHI (Figure 8). Overall, our study demonstrates that CRISPR-Cas is a barrier to antibiotic resistance dissemination in the manure environment and identifies key areas for future study of CRISPR-Cas efficacy in the agricultural setting.

## ACKNOWLEDGEMENTS

We thank US Department of Agriculture for their collaboration and members of the Palmer lab for their constant guidance and feedback. We are grateful to Dr. Robert Dungan (USDA ARS, Kimberly, Idaho) for the dairy manure sample that was used for the experiments. This study was funded by the USDA-ARS Office of National Programs (Project No. 58-3042-8-012) and administered via National Program 202 Soil and Air. This work was also supported by R01AI116610 from the National Institutes of Health and the Cecil H. and Ida Green Chair in Systems Biology Science to K.P.

